## Supplementary figures and images for "An immunohistochemical atlas of necroptotic pathway expression"

### Fig. EV1

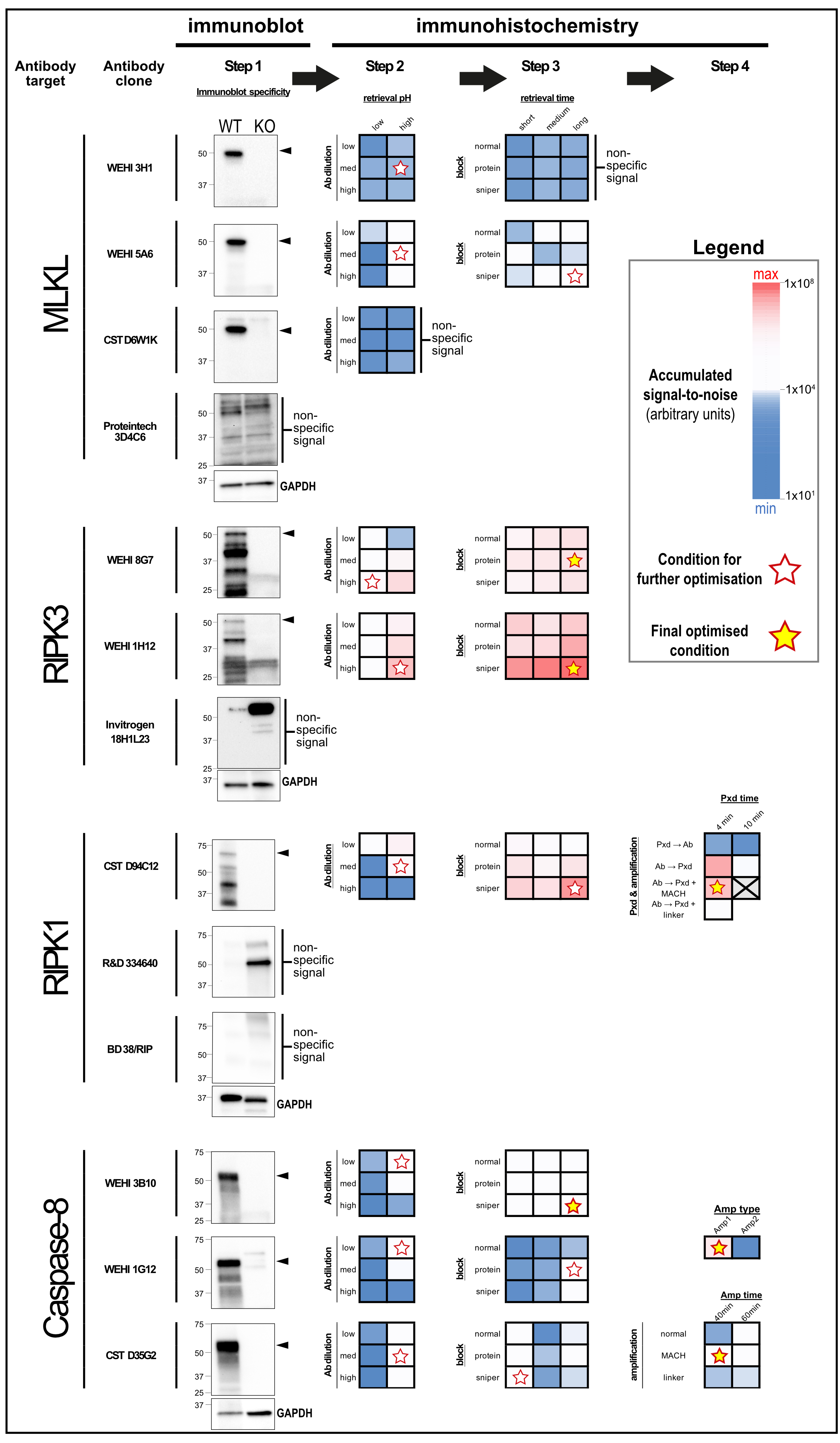

### Fig. EV2

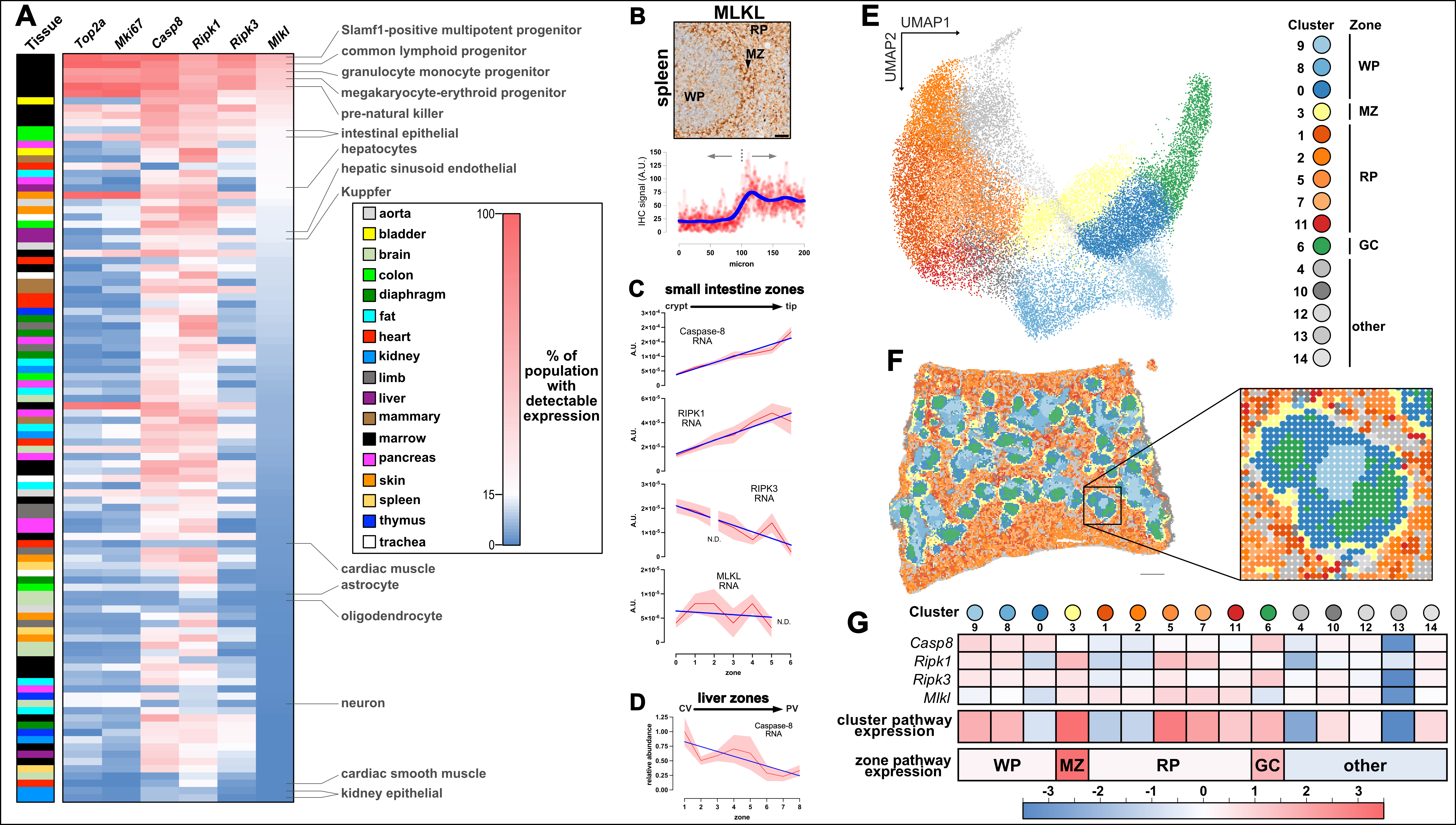

### Fig. EV3

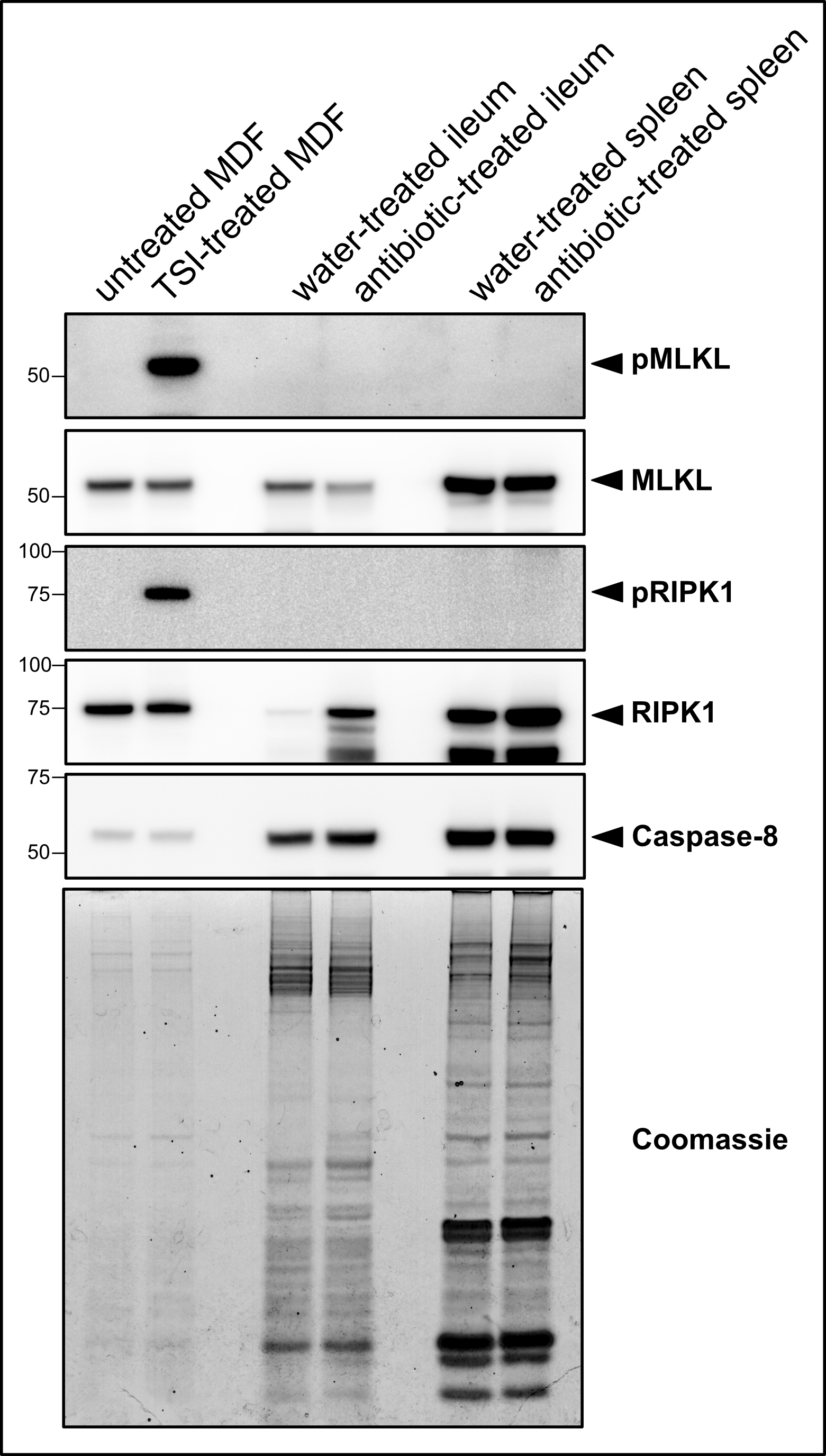

### Fig. EV4

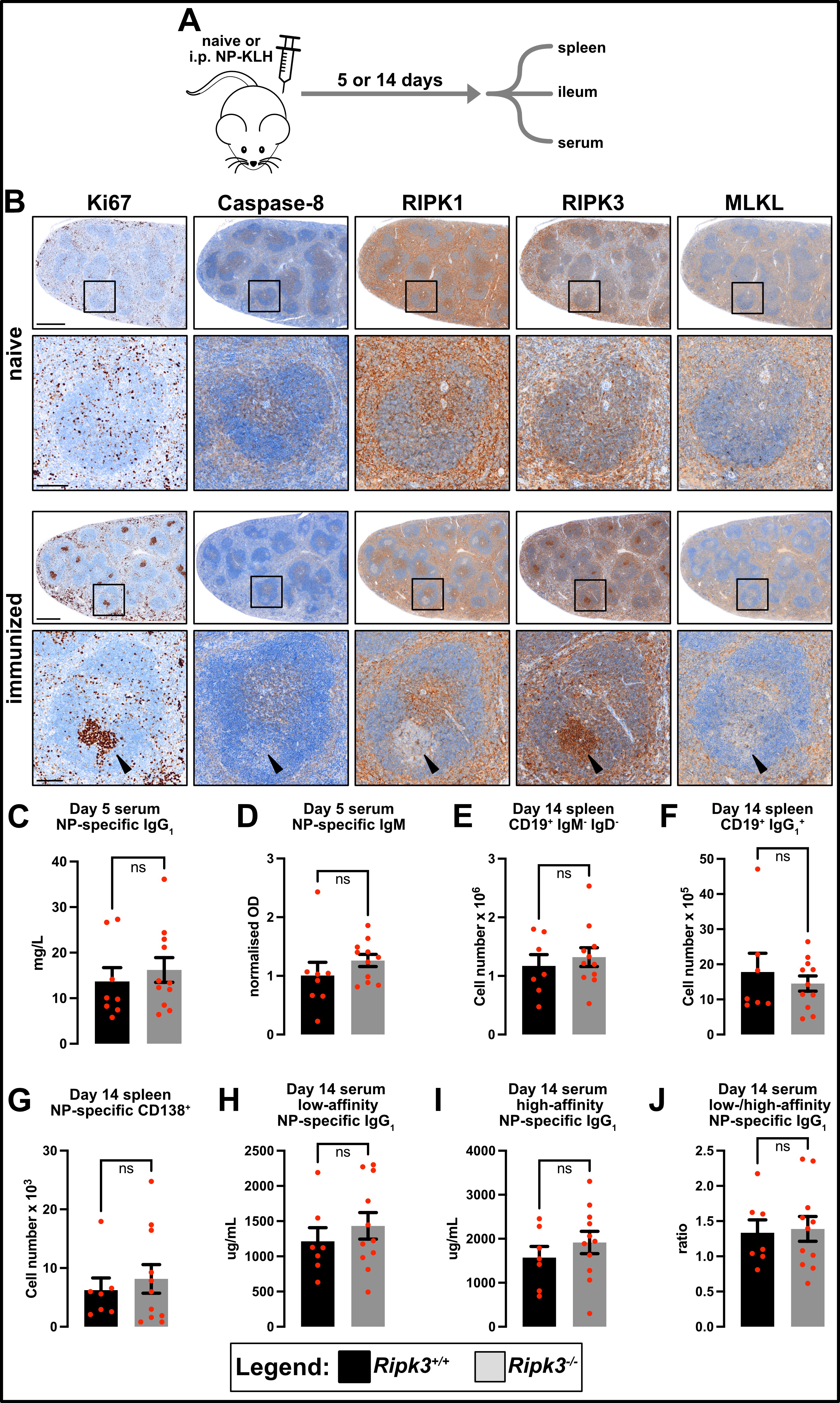

### Fig. EV5

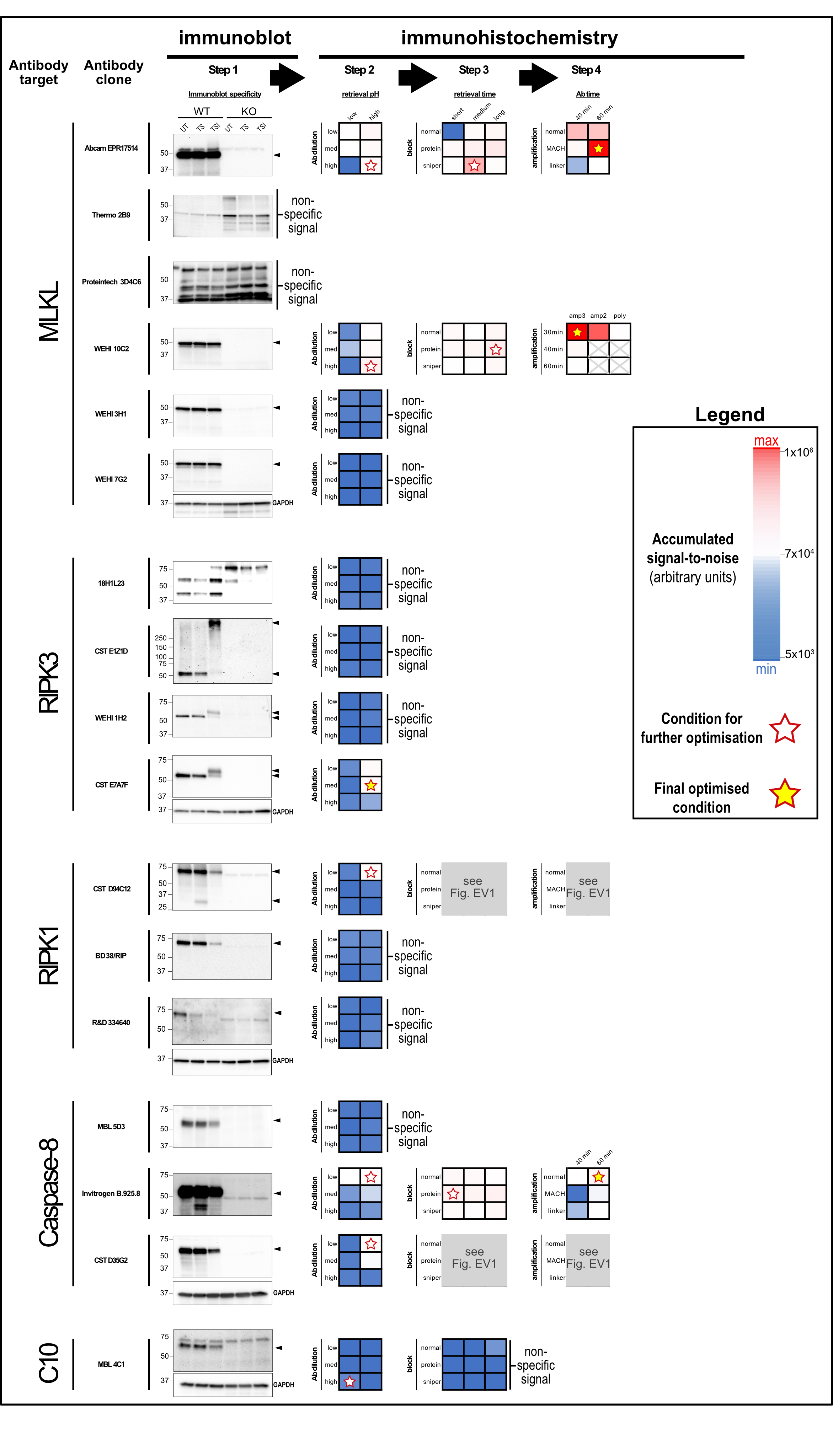

### Fig. EV6

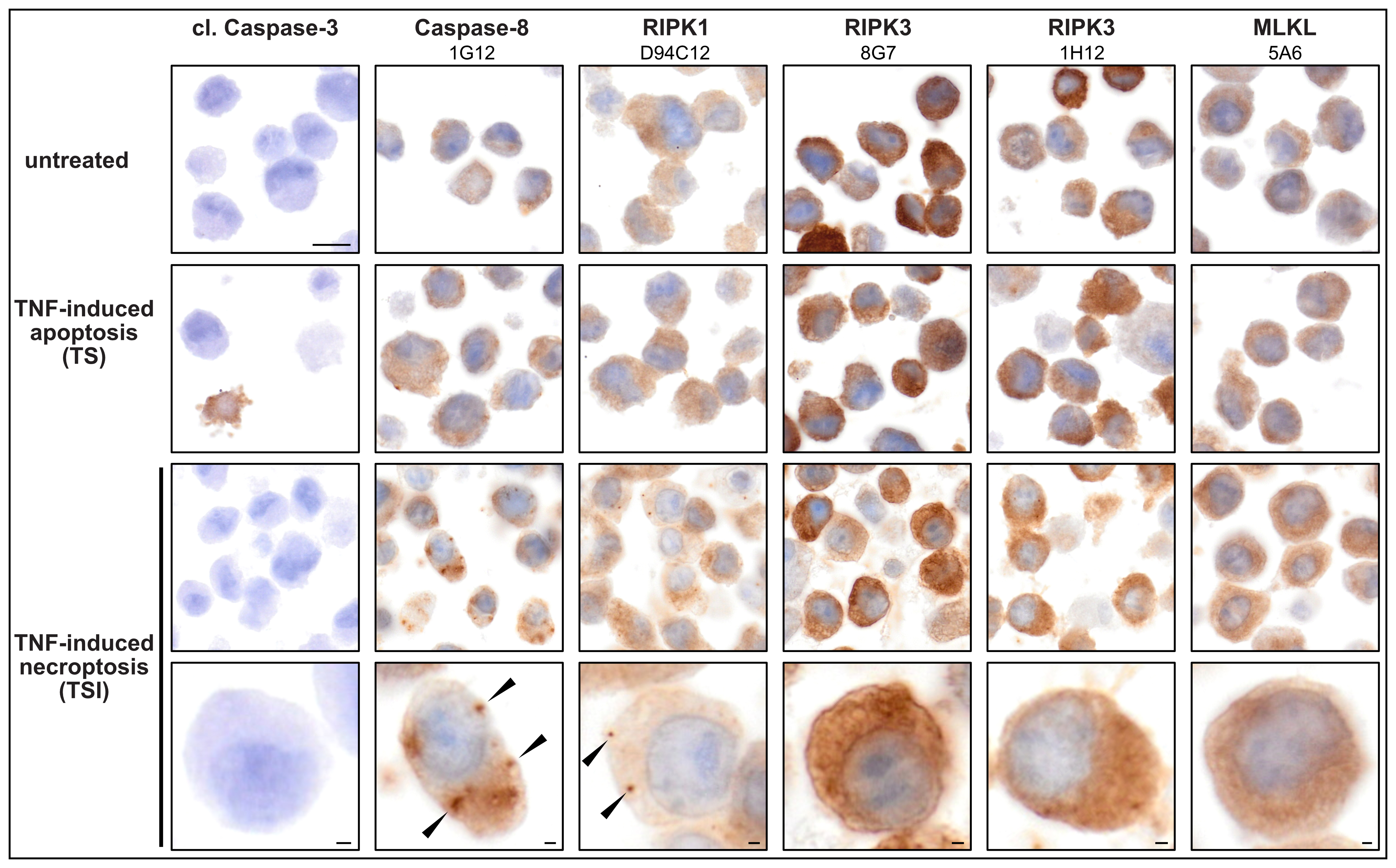

### Fig. EV7

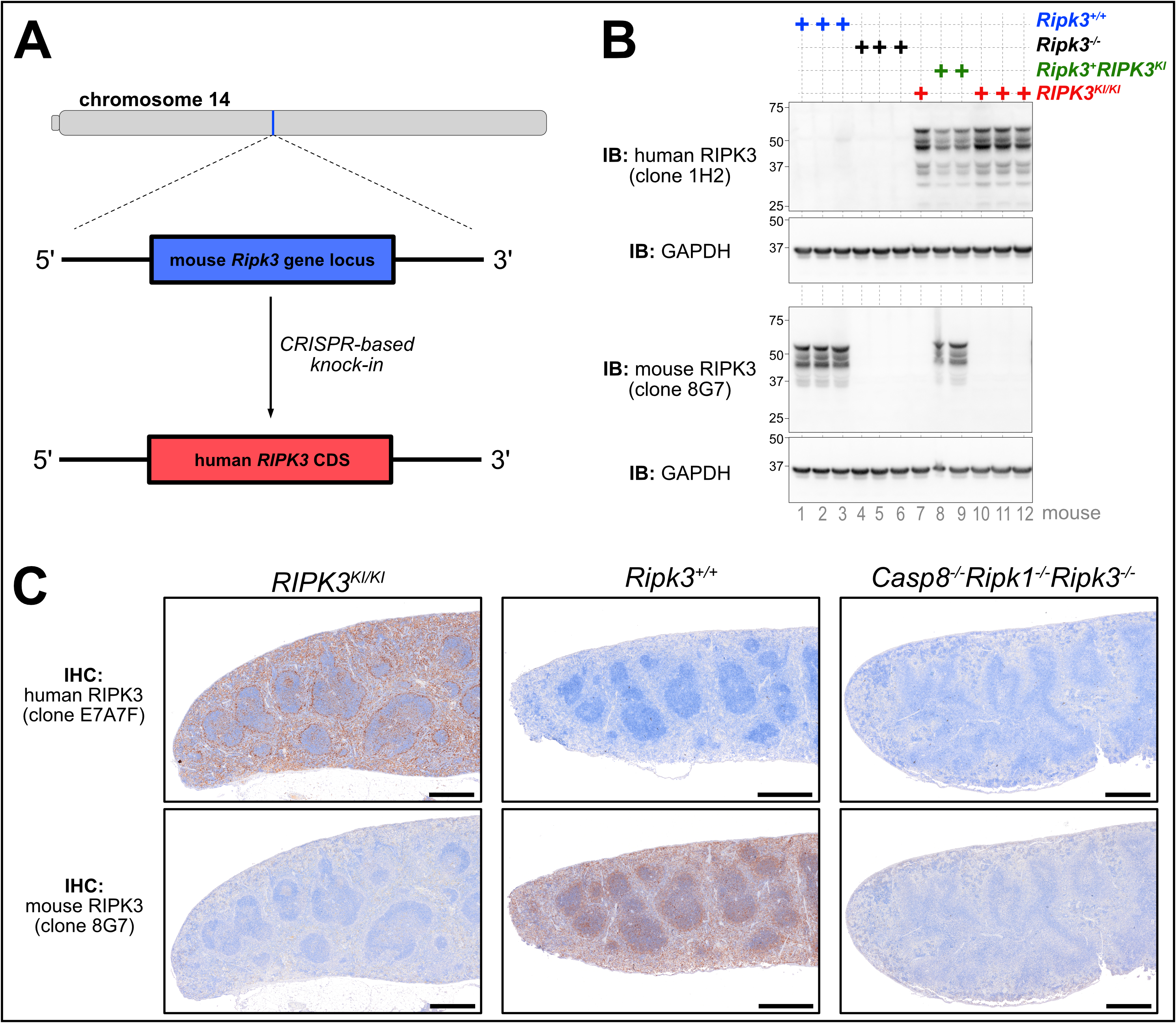

### Fig. EV8

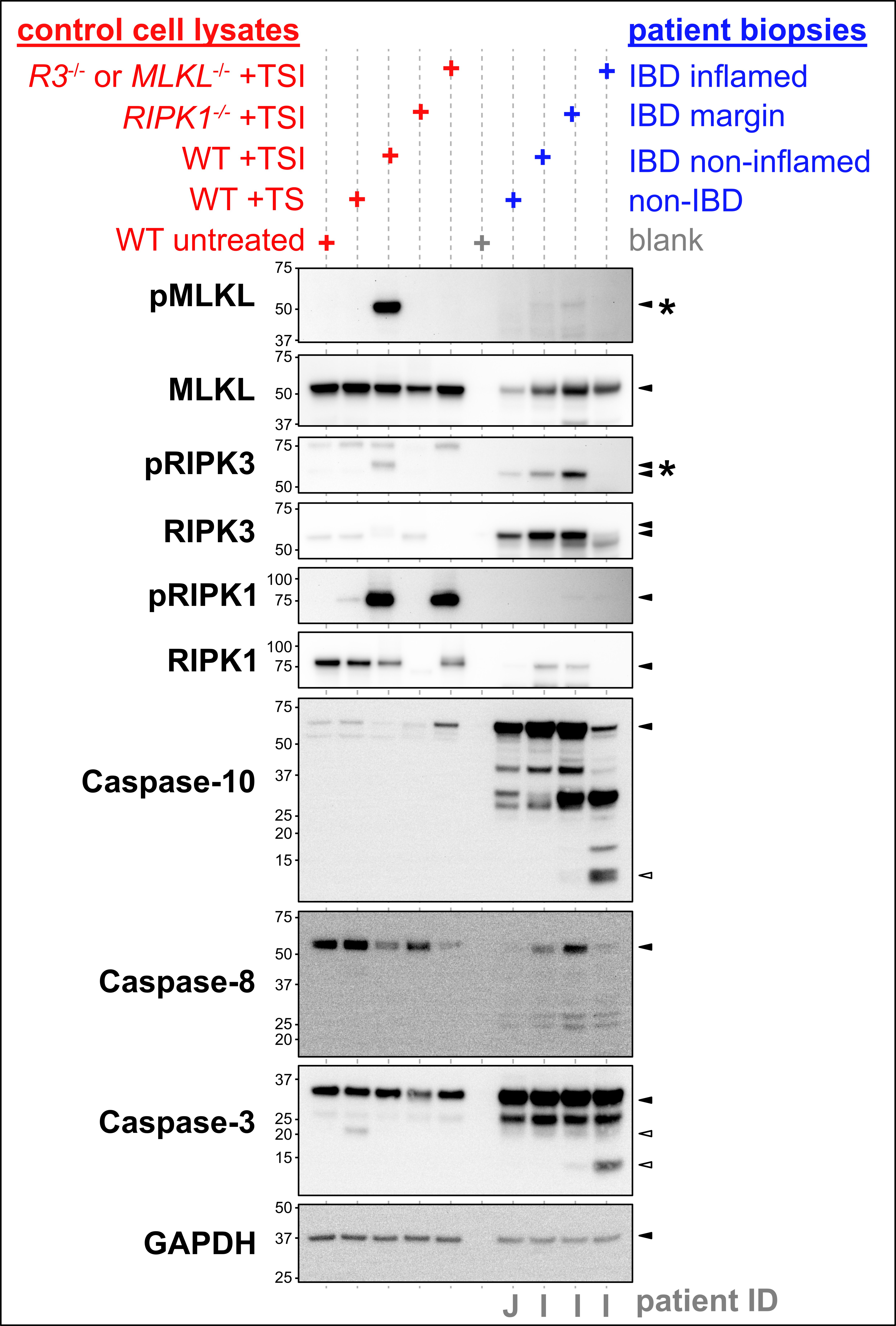

### Fig. EV9

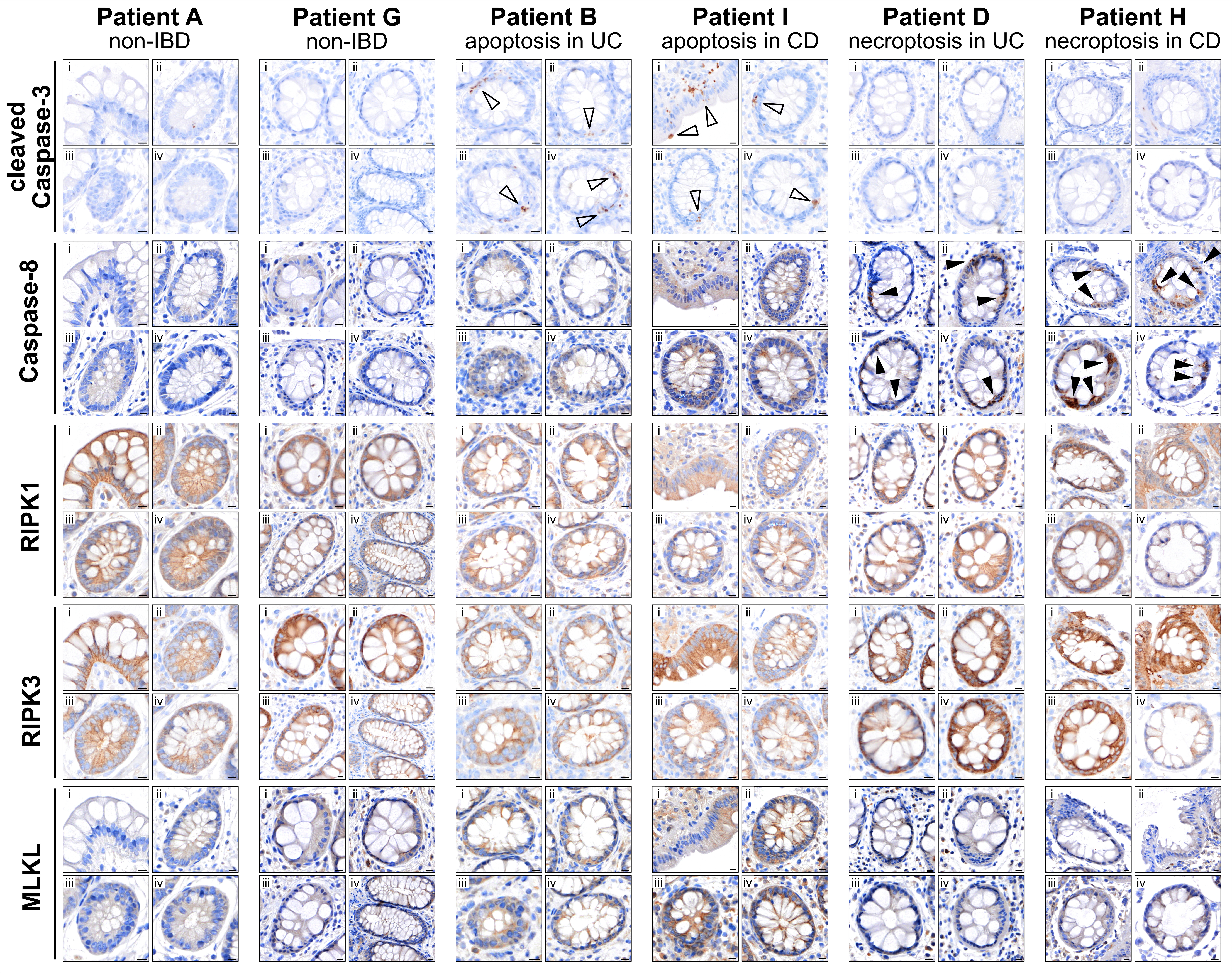

### Fig. EV10

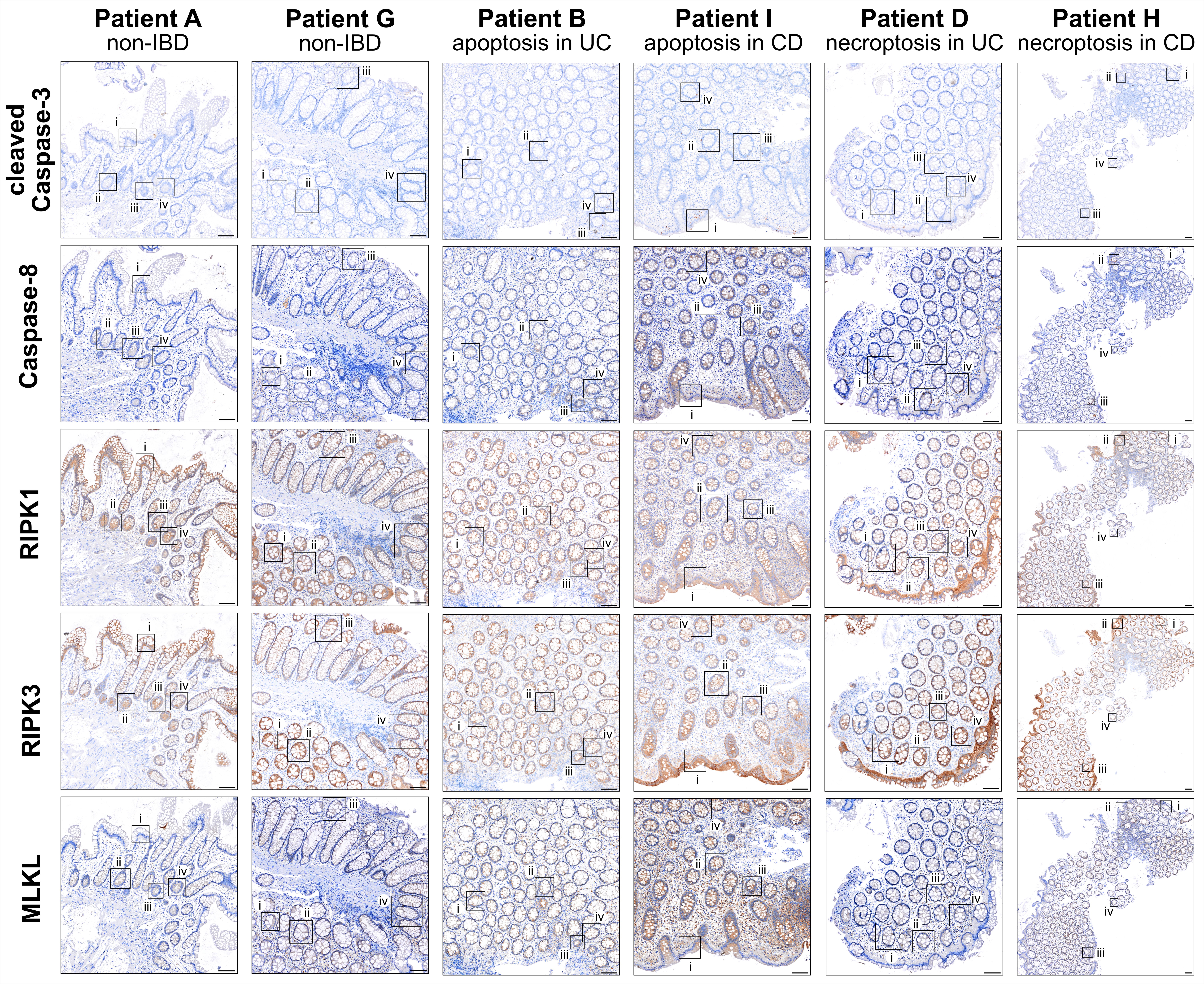

### Supplementary Table 1

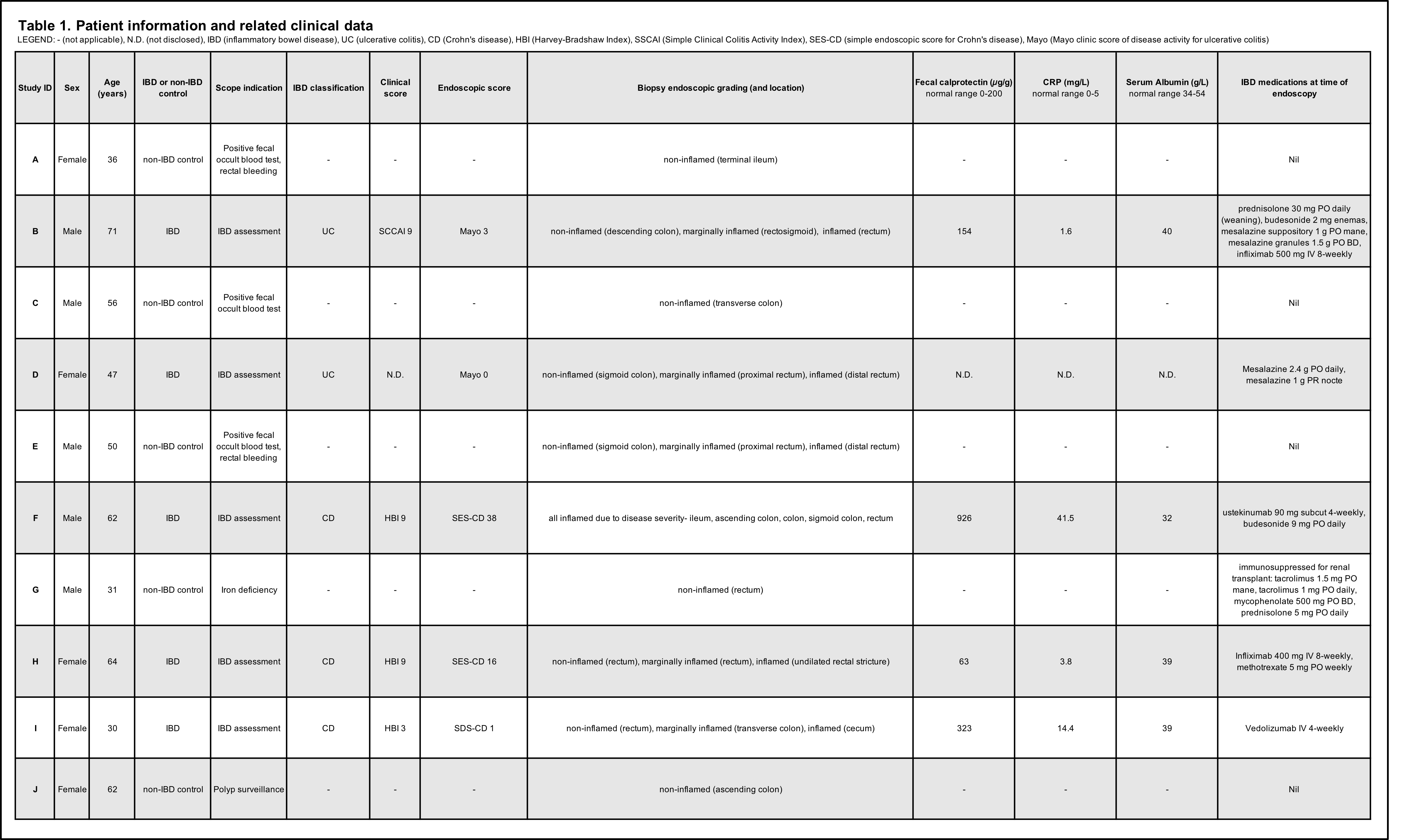
